# Regulation of meiotic progression by Sertoli-cell androgen signaling

**DOI:** 10.1101/2020.05.26.117093

**Authors:** Hailey Larose, Travis Kent, Qianyi Ma, Adrienne Niederriter Shami, Nadia Harerimana, Jun Z. Li, Saher Sue Hammoud, Mary Ann Handel

## Abstract

Androgen receptor (AR) signaling in Sertoli cells is known to be important for germ-cell progression through meiosis, but the extent to which androgens indirectly regulates specific meiosis stages is not known. Here, we combine synchronization of spermatogenesis, cytological analyses and single-cell RNAseq (scRNAseq) in the <u>S</u>ertoli <u>c</u>ell <u>a</u>ndrogen <u>r</u>eceptor <u>k</u>nock<u>o</u>ut (SCARKO) mutant and control mice, and demonstrate that SCARKO mutant spermatocytes exhibited normal expression and localization of key protein markers of meiotic prophase events, indicating that initiation of meiotic prophase is not androgen dependent. However, spermatocytes from SCARKO testes failed to acquire competence for the meiotic division phase. ScRNAseq analysis of wild type and SCARKO mutant testes revealed a molecular transcriptomic block in an early meiotic prophase state (leptotene/zygotene) in mutant germ cells, and identified several misregulated genes in SCARKO Sertoli cells, many of which have been previously implicated in male infertility. Together, our coordinated cytological and single-cell RNAseq analyses identified germ-cell intrinsic and extrinsic genes responsive to Sertoli-cell androgen signaling that promotes cellular states permissive for the meiotic division phase.

## Introduction

Mammalian germ cells are in close contact with multiple testicular somatic cells that provide instructive cues for their development and differentiation. In the testes, Sertoli cells, are the only niche cells within the seminiferous epithelium that are in contact with germ cell populations to support all stages of their development, including meiosis which ultimately leads to production of haploid gametes. Previous studies have shown that androgens, and specifically androgen receptor (AR) signaling in Sertoli-cells, is one mechanism by which meiotic progress is regulated, but exactly how AR signaling in neighboring Sertoli cells promotes the events of meiosis is not well understood.

During the substages of the first meiotic prophase (leptonema, zygonema, pachynema and diplonema), homologous chromosomes line up to form diploid pairs (synapsis), and undergo recombination. At the end of the meiotic prophase, spermatocytes rapidly undergo the first reductive division followed by the second equational division to generate 4 haploid gametes (described in detail in Bolcun-Filas and Handel, 2018). Multiple kinases have been implicated to regulate the onset of the first meiotic division, termed the G2-MI transition, but the mechanism by which it is initiated have been difficult to delineate (Sun and Handel, 2008; Jordan *et al.*, 2012). However, we know that spermatocytes acquire competency for the G2-MI transition several days before the division occurs. This was revealed in early work (Cobb *et al.*, 1999) in which okadaic acid (OA), a pleitropic phosphatase inhibitor used in a variety of systems to prompt cell division-related condensation of chromosomes, was found to trigger precocious induction of the G2-MI transition in male germ cells. Competency to undergo this response to OA is acquired concomitently with the appearance of the H1t histone variant, formally known as H1F6 (Cobb *et al.*, 1999), but the H1t histone variant is not required for the germ cell to enter the meiotic division phase (Lin *et al.*, 2000).

Progress of meiotic prophase and acquisition of meiotic competence is typically perceived as germ cell-intrinsic processes, but emerging molecular profiling studies suggest that neighboring testicular somatic cells, and in particular Sertoli cells, may directly or indirectly control these meiotic events. Interestingly, Sertoli cells undergo distinct molecular and metabolic changes across the seminiferous tubule. In adult mammals, the seminiferous epithelium regularly cycles through distinct stages, each one of which is morphologically defined by recurring germ cell states (e.g., characteristic associations of spermatogonial populations, meiotic prophase cells, round/elongating spermatids, etc.), as well as distinct molecular and metabolic Sertoli cell states (Chen *et al.*, 2018; Green *et al.*, 2018). These histological associations define the stages (12 in mice) of the seminiferous epithelium cycle, setting both the environment and the pace within which germ cell differentiation unfolds (Kent *et al.*, 2019).

One required and dynamically regulated signaling pathway across the seminiferous epithelial stages is androgen signaling. Genetic loss-of-function experiments show that global knockout (KO) of the *Ar* gene is required for testis development and spermatogenesis (O’Hara and Smith, 2015), although the germ cells themselves do not require cell-autonomous expression of androgen receptors (Johnston *et al.*, 2001). AR is expressed by many somatic cell types in the testis, but *A*r transcript and protein levels in Sertoli cells peak at Stages VI-VII - coincident with differentiating spermatogonia committing to meiotic prophase entry(O’Hara and Smith, 2015) suggesting that Sertoli cell AR signaling may mediate stage-specific roles in meiosis. Indeed, in a conditional <u>S</u>ertoli-<u>c</u>ell only *<u>Ar</u>*<u>k</u>nock<u>o</u>ut referred to as the SCARKO mouse model spermatogenesis is arrested in meiosis (De Gendt *et al.*, 2004), but there are some differences among reports as to which specific events of meiosis are affected (Chen *et al.*, 2016).

In our work, we asked if androgen signaling is required for acquisition of competence for meiotic division and, if so, how androgen shapes the transcriptional landscape of spermatocytes to instruct progress through meiotic prophase. As a result of asynchronous and iterative waves of spermatogenesis progress across the entire seminiferous epithelium, leading to multiple germ cell populations within a given cross section of the testis, it proved difficult to parse out interactions among major events of meiosis and their stage specificity in early work. More recently, methods were developed to manipulate availability of RA in the mouse testis to precisely synchronize the onset of spermatogenesis, so that only 1 to 3 temporally adjacent stages are represented at any moment (Hogarth *et al.*, 2013, 2015; Griswold, 2016).Thus, it is possible to apply this methodology to relate stage-specific Sertoli cell signaling events to the spermatogenic, and more specifically, meiotic, events they control. Using stage-synchronized testes samples (**Figure 1; Supplemental Table 1**) from the SCARKO mice, we here investigated the cytological hallmarks of meiotic prophase and the temporal acquisition of division-phase competence. We also performed parallel single-cell transcriptomic (scRNAseq) analyses to determine androgen-regulated transcriptional markers of meiotic progress.

**Figure 1.**
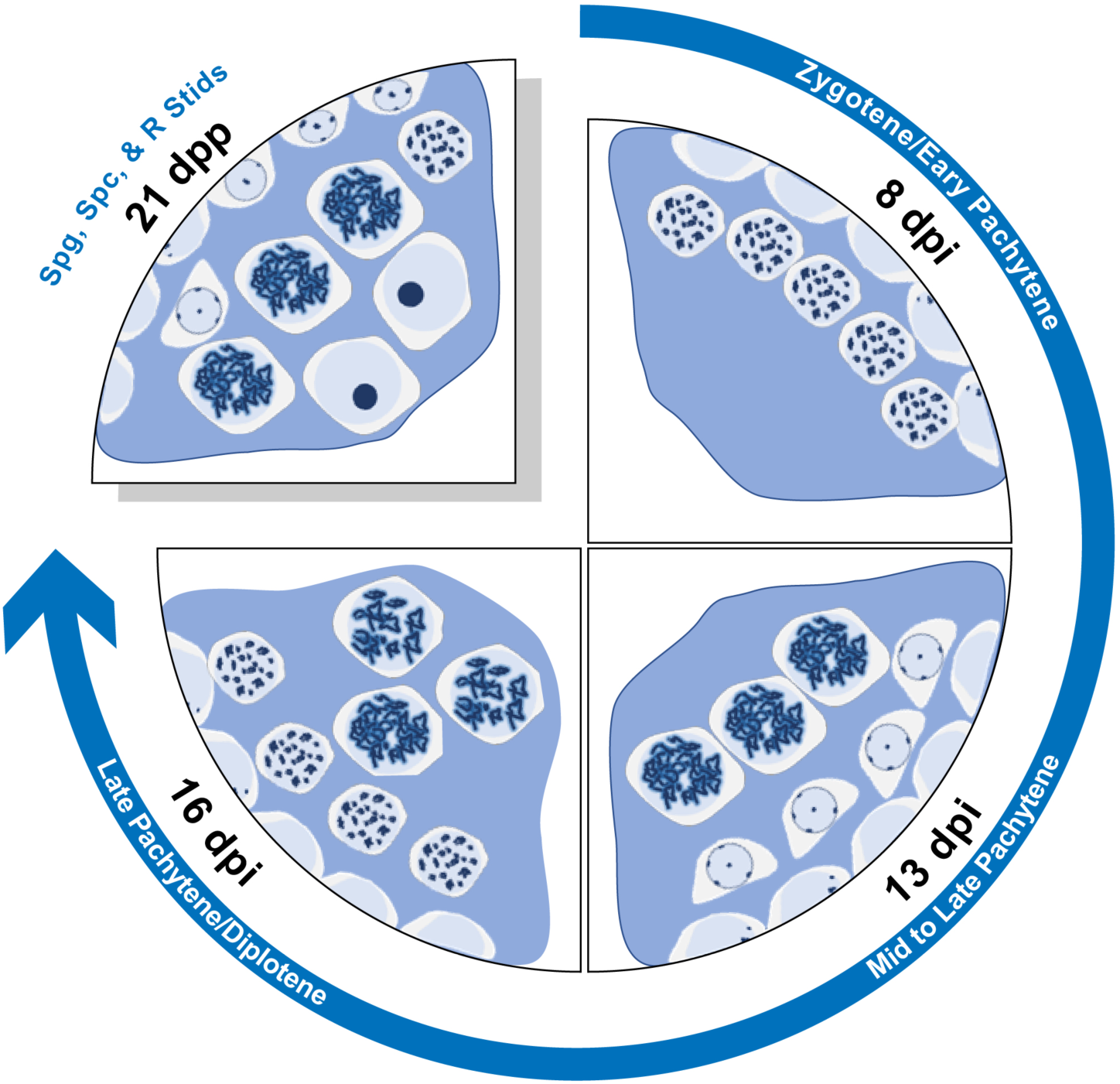
Experimental strategy and source of cells for scRNAseq. In this schematic of seminiferous tubule cross-sections, the first three quadrants, with the blue arrow, illustrate the cell populations retrieved at 8, 13, and 16 days after injection (dpi) of retinoic acid to synchronize spermatogenesis. The fourth quadrant illustrates the germ-cell composition of seminiferous tubules retrieved at 21 days of age (21 dpp), with no manipulation of retinoic acid.

We confirmed that Sertoli-cell AR signaling is required for the overall survival of meiotic prophase spermatocytes, while in the absence of AR, germ cells are progressively lost. Among surviving spermatocytes that enter meiotic prophase in SCARKO mice testes, chromosome synapsis and recombination occurred normally, suggesting that early meiotic prophase events are not dependent on Sertoli-cell AR signaling. However, we discovered that Sertoli cell AR signaling is required for spermatocytes to acquire competence to enter the meiotic division phase, explaining the observed cytological arrest in spermatocytes of SCARKO testes. To achieve a comprehensive and unbiased molecular understanding of the SCARKO phenotype, we leveraged our testicular germ cell synchronization (**Figure 1; Supplemental Table 1**) and single-cell transcriptomics analysis, which enabled us to pinpoint a set of spermatocyte-expressed genes regulated by AR signalling in a non-cell-autonomous manner. Many of the genes have likely or known post-meiotic roles, suggesting that this gene set may function to license spermatocytes for progression towards spermiogenesis. Overall, these findings reveal the molecular machinery by which androgen signaling in Sertoli cells creates a permissive environment for germ cell progression through meiosis and gamete production.

## Results

### Sertoli cell-expressed androgen receptor is functionally required to maintain germ cell numbers

Consistent with previous reports on additional SCARKO models (De Gendt *et al.*, 2004); (Chang *et al.*, 2004), our histological analyses of testes from 21 days post partum (dpp) SCARKO males (**Supplemental Table 1**) revealed both loss of meiotic germ cells and failure of surviving spermatocytes to progress. Most spermatocytes appeared arrested at early to mid-meiotic prophase I, with very few completing the first meiotic division (**Figure 2A).** To assess the extent of apoptosis, we synchronized seminiferous tubules in experimental animals and conducted TUNEL analysis on wild-type and synchronized SCARKO testes collected at 8, 10, 12, 14, and 16 days post-RA injection (dpi) (**Figure 2B; Supplemental Table 1**). As expected, histological testes sections from wild-type animals were enriched for germ cells at late leptotene/early zygotene (8 dpi), late zygotene/early pachytene (10 dpi), early/mid pachytene (12 dpi), mid/late pachytene (14 dpi), and late pachynema/diplotene (16 dpi) stages (**Supplemental Table 1**), however, the percentage of apoptotic cells per tubule in SCARKO testes was significantly increased in at all stages except preleptonema (**Figure 2B**). These histological observations are consistent with both progressive germ-cell loss due to apoptosis and failure to differentiate beyond meiotic prophase in the absence of androgen signaling from Sertoli cells.

**Figure 2.**
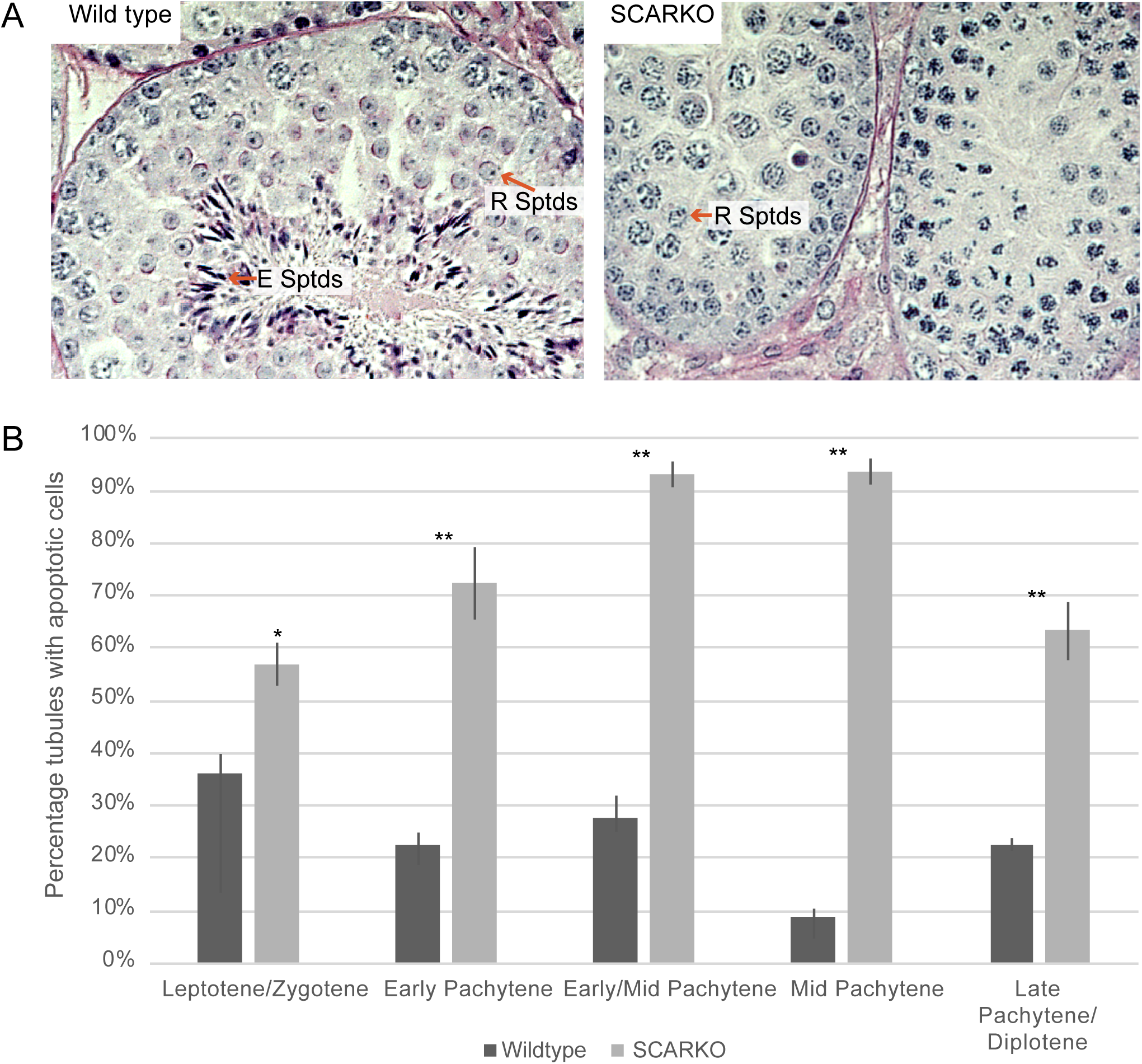
SCARKO testes exhibit germ-cell arrest and progressive loss. **A**. Cross sections of wild type (L) and SCARKO (R) adult testes. Arrows and arrowheads represent round and elongated spermatids respectively. **B.** Percentage of tubules with TUNEL-positive cells from testes of both wild type and SCARKO males at 8, 10, 12, 14, and 16 DPI (days post RA injection).

### Spermatocytes in SCARKO mice enter prophase, but fail to acquire competence for initating meiotic division

Given the almost complete absence of post-meiotic germ cells in SCARKO testes, we determined whether classical protein markers for meiotic prophase milestones were expressed in spermatocytes. We assessed markers for chromosome pairing and synapsis, proteins involved in meiotic homologous recombination, and the differentiation marker (histone H1t), whose expression coincides with onset of pachytene and acquisition of competence for meiotic division (Inselman *et al.*, 2003, Cobb *et al.*, 1999). To assess homologous chromosome pairing and the synaptonemal complex (SC) integrity, we used antibodies recognizing the SC-associated proteins SYCP3 and SYCP1 (marking the lateral and central element established at synapsis, respectively). Spermatocytes from both wild-type and SCARKO testes appeared to be capable of assembling normal SCs, as indicated by labeling for SYCP3 and SYCP1 (**Figure 3A,C**). In early stages of meiotic prophase, phosphorylated histone H2AX (γH2AX) is dispersed throughout the chromatin, indicating formation of recombination-related double-strand breaks (DSBs). In the pachytene stage, γH2AX becomes restricted to the XY body - indicating that most DSBs have been repaired. When examining these events in both the synchronized or nonsynchronized wild-type and SCARKO spermatocytes we found that γH2AX staining patterns were generally similar, although some abnormalities were observed in spermatocytes from SCARKO mutants (**Figure 3A,B**). Similarily, MLH1 distribution, which corresponds to the number and distribution of chiasmata, was comparable in wild-type and SCARKO spermatocytes **(Figure 3D**). Finally, we assessed whether the germ cells progress through pachynema by staning for histone H1t (Inselman *et al.*, 2003). However, because germ cells in SCARKO testes gradually become depleted due to apoptosis, we analyzed germ cells at the same developmental stages from synchronized wild-type and SCARKO testes, and found no differences in the number of H1t-positive spermatocytes (**Figure 3E-F**). This observation indicates that mid-pachytene expression and localization of H1t on chromsome is not regulated by Sertoli-cell AR signaling. Taken together, our analysis of meiotic prophase markers suggest that loss of AR in Sertoli cells does not impede chromosome synapsis, recombination events, or accumulation of histone H1t in spermatocytes that escape apoptosis.

**Figure 3.**
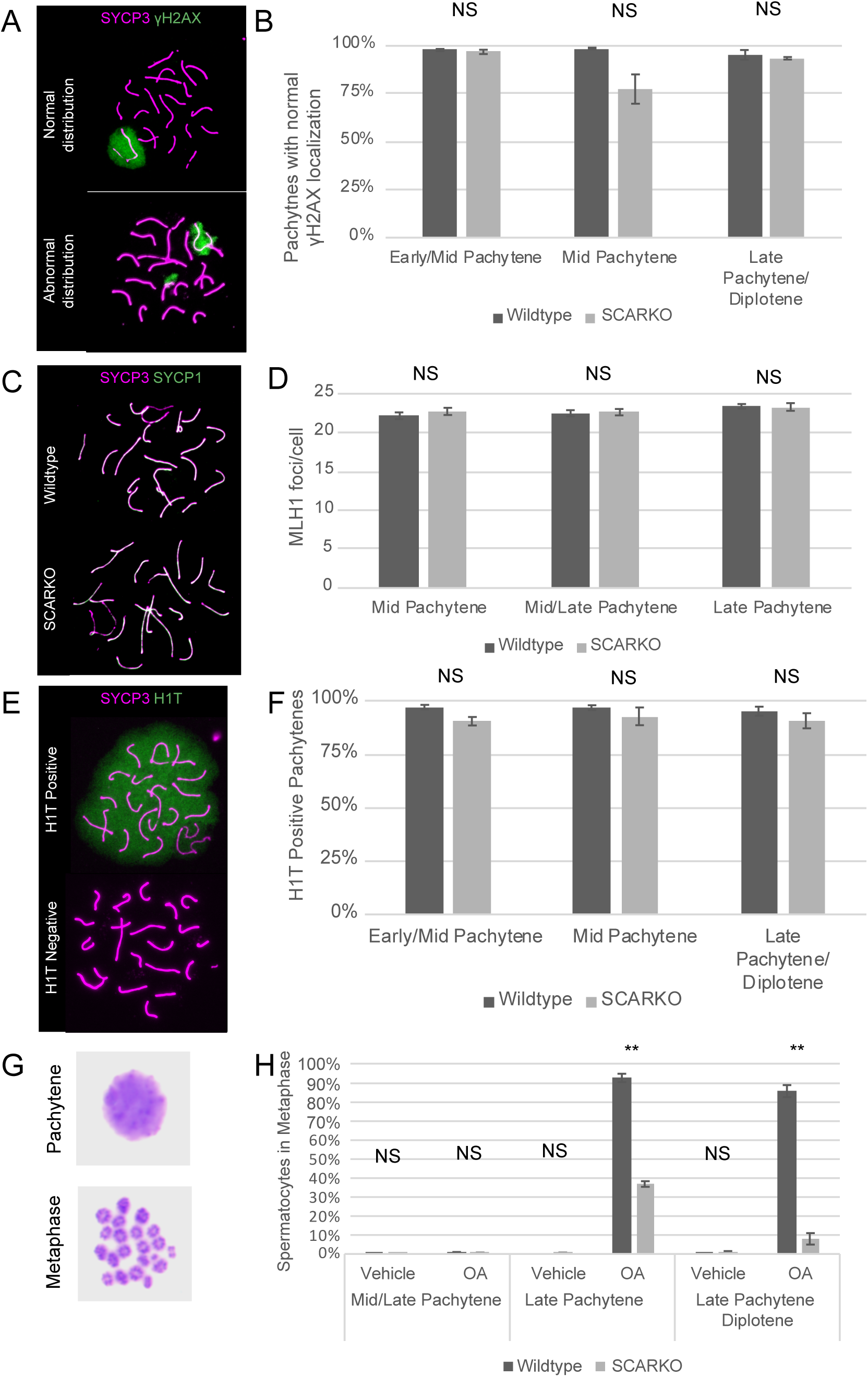
Many features of meiosis are normal in SCARKO testes, but spermatocytes do not acquire competence for the OA-induced meiotic division phase. **A.** At the pachytene sub-stage of meiotic prophase, SCARKO spermatocytes exhibit both normal distribution of γH2AX, restricted to the XY body, and abnormal patterns with γH2AX also in the autosomal domain. **B.** Distribution of γH2AX scored in pachytene spermatocytes isolated from wild type and SCARKO synchronized testes, with the X axis indicating the stages of pachytene cells. **C.** This panel shows representative SYCP1 (green) and SYCP3 (magenta) localization in pachytene spermatocytes isolated from wild type (top) and SCARKO (bottom) testes. **D.** Quantification of MLH1 foci in pachytene spermatocytes isolated from wild type and SCARKO testes. The x-axis indicates the stages of pachytene cells isolated from synchronized testis. **E.** This panel shows H1T-positive and -negative pachytene spermatocytes from wild type and SACRKO testes. **F.** This graph demonstrates similar frequency of H1T-positive spermatocytes from wild type and SCARKO synchronized testes. The x-axis indicates the stages of pachytene cells isolated from synchronized testis. **G.** This illustrates typical Giemsa-stained pachytene (top) and metaphase (bottom) spermatocytes after OA treatment. **H**. The graph presents the percentage of metaphase cells among both wild type and SCARKO spermatocytes after treatment with OA. The x-axis indicates both the treatment the cells were exposed to, as well as the prophase substage of the most advanced spermatocytes at the time of isolation. (NS – Not significant; ** - P<0.01)

These observations raise a fundamental question: Why do germ cells that progress to mid to late meiotic prophase in SCARKO testes fail to undergo meiotic division? To answer this question, we treated spermatocytes with the phosphatase inhibitor OA, a regime that induces chromosome condensation and entry into metaphase by meiotically competent spermatocytes (Cobb *et al.*, 1999). Testes were isolated at 14, 15, and 16 dpi, when spermatocytes correspond to those in seminiferous epithelial Stages VII-VIII, VIII-IX, and X-XI, respectively, which are enriched for spermatocytes at mid-to-late pachynema. As expected, OA treatment resulted in spermatocytes in a pachytene configuration, or, from metaphase-competent spermatocytes, with condensed metaphase bivalents (**Figure 3G**). We did not observe any OA-induced condensation of meiotic bivalents at 14 dpi in spermatocytes from either SCARKO or wild-type testes **(Figure 3H**). In contrast, only 5-25% of pachytene spermatocytes from synchronized SCARKO mutant mice collected at 15 and 16 dpi were competent to condense chromosomes and enter metaphase, compared with ∼90-95% from wild-type mice (**Figure 3H**). This analysis reveals that although SCARKO spermatocytes are at comparable cytological state as wild-type cells, they have not acquired the inherent competence to respond to OA. This functional difference may stem from AR signaling-dependent molecular circuitry governing acquisition of competency.

### scRNAseq reveals early meiotic arrest of spermatocytes in SCARKO mutants

We next set out to investigate how absence of AR signaling from surrounding Sertoli cells impacts on the transcriptional landscape underlying acquisition of meiotic competency.To this end, we performed single-cell RNA sequencing (scRNAseq) on whole testis cell suspensions from 21 dpp wild-type and SCARKO males, an age in which all germ cell stages of interest are present (**Figure 1; Supplemental Table 1**), and from synchronized testes. To enrich for germ cells, we collected tissue at three timepoints post RA injection corresponding to key meiotic prophase transitions including zygotene to early pachytene (8dpi), mid-pachytene (13dpi), and late pachytene to diplotene transition (16dpi) (**Figure 1; Supplemental Table 1**). From each sample, we collected an average of ∼3,300 cells, with a total of 26,500 sequenced cells. After quality control (QC) filtering, a total of 21,314 cells were retained across all datasets with an average of ∼1,800 detected genes per cell (see Methods). Systematic assessment of batches confirmed cluster reproducibility and revealed high cluster-cluster correlations across biological timepoints, and consistent with the expected outcome of the synchronization treatment, a few germ cell clusters detected unevenly across dataset (**Figure S1A,B**). Given the high concordance across datasets, we merged the eight datasets (both wild type and SCARKO) for unsupervised clustering. Within the compiled dataset of ∼21,000 cells we identified seven clusters **(Figure 4A**), with each molecular cluster consisting of cells from multiple datasets (**Figures 4B and S1B**). Using the expression pattern of previously known cell type-specific markers (Chen *et al.*, 2018; Green *et al.*, 2018; Jung *et al.*, 2019), we identified the major expected germ-cell types, including spermatogonia and spermatocytes, as well as five somatic cell types (**Figure 4C, Supplemental Table S2).**

**Figure 4:**
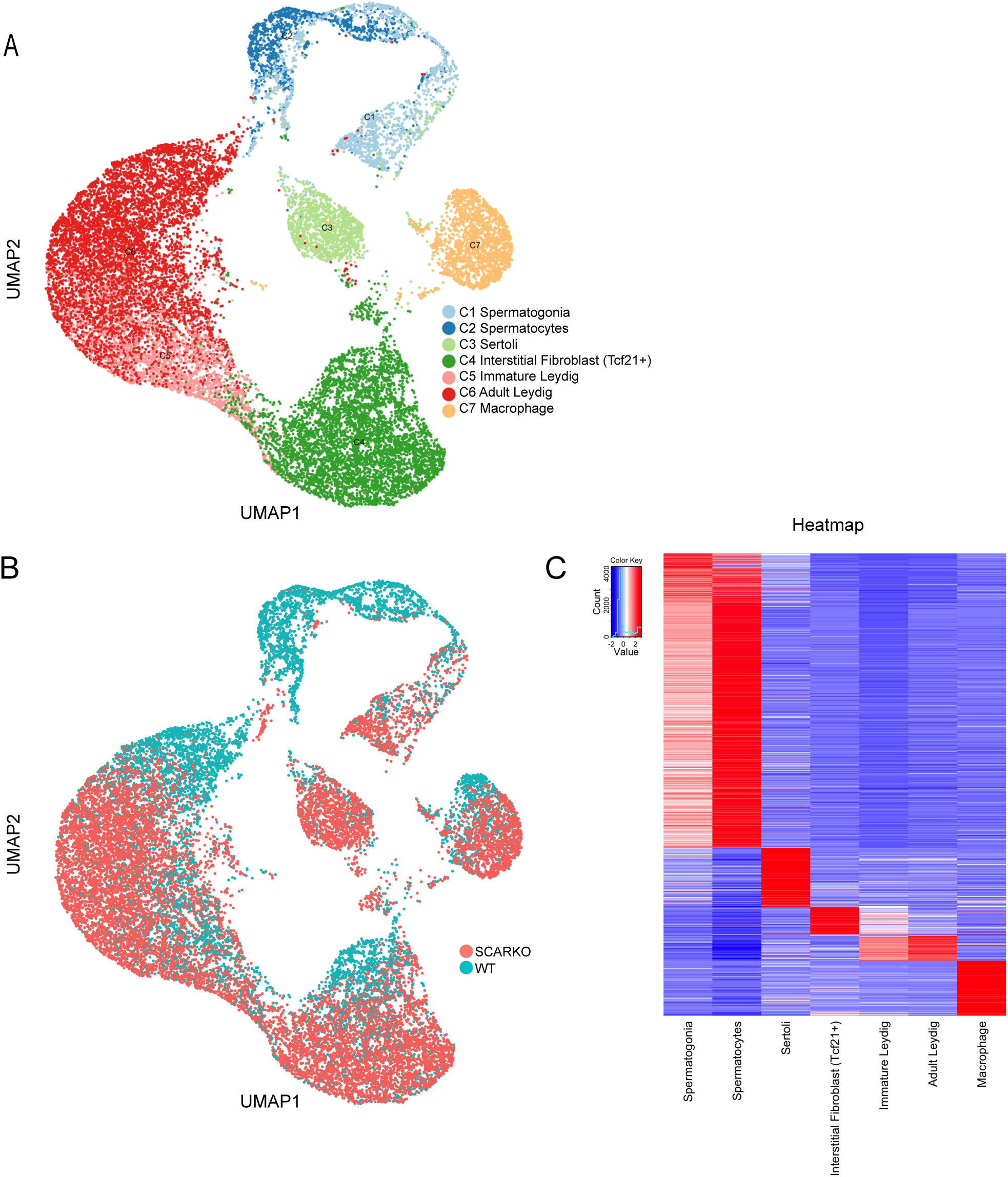
Overview of Major Cell Types and Cellular Attributes Inferred from Single-Cell RNA-Seq Analyses of wild-type and SCARKO mouse testes. A) Cellular heterogeneity at the highest levels regardless of genotype or time-point of collection. Principal component analysis of all 21,000 cells post-quality control (QC) filter reveals 7 clusters, corresponding to two germ cell populations and five somatic cell types. Data visualized in UMAP space. B) Visualization of all 21,000 cells post-quality control (QC) filter in UMAP space, overlaid by genotype contribution. C) Marker gene Heatmap of top-expressed markers for the 7 cell types. Note – in the heatmap each marker gene is standardized over the 7 cluster centroids and ordered by cell type.

To identify specific stages of arrest and transcriptional changes that differentiate germ cells in SCARKO from those in wild-type testes, we re-clustered only the germ-cell populations from all eight datasets, and obtained seven major clusters (**Figure 5A; Supplemental Table S3**). As inferred from expression of previously known cell-specific marker genes, these clusters correspond to undifferentiated spermatogonia (C1), differentiating spermatogonia (C2), leptotene/zygotene spermatocytes (C3), early-mid pachytene spermatocytes (C4), late pachytene spermatocytes (C5), diplotene spermatocytes (C6) and early round spermatids (C7) (**Figure 5A,B, Supplemental Table S3**) (Chen *et al.*, 2018; Jung *et al.*, 2019). More mature germ cells were not expected, given collection from juvenile testes, **Figure 1**. Although several markers are shared between undifferentiated and differentiated spermatogonia, some were more enriched in either undifferentiated spermatogonia (C1) or differentiating spermatogonia (C2; e.g. *Gfra1* and *Zbtb16* in C1 and *Stra8, Dmrt1* in C2; see all markers in **Supplemental Table S3**). In the leptotene/zygotene stage (C3) cluster, we found that the spermatogonial cell transcriptional pattern is no longer dominant, and that instead a distinct set of genes is upregulated (**Figure 5B and Supplemental Table S3**). Among cells in the early-mid pachytene cluster (C4), further changes were observed in the transcriptome, with the spermatogonial transcriptional pattern being absent and many genes distinctively upregulated, in a pattern continuing from pachytene through the diplotene transition, C5-7 **(Figure 5B, Supplemental Table S2).**

**Figure 5:**
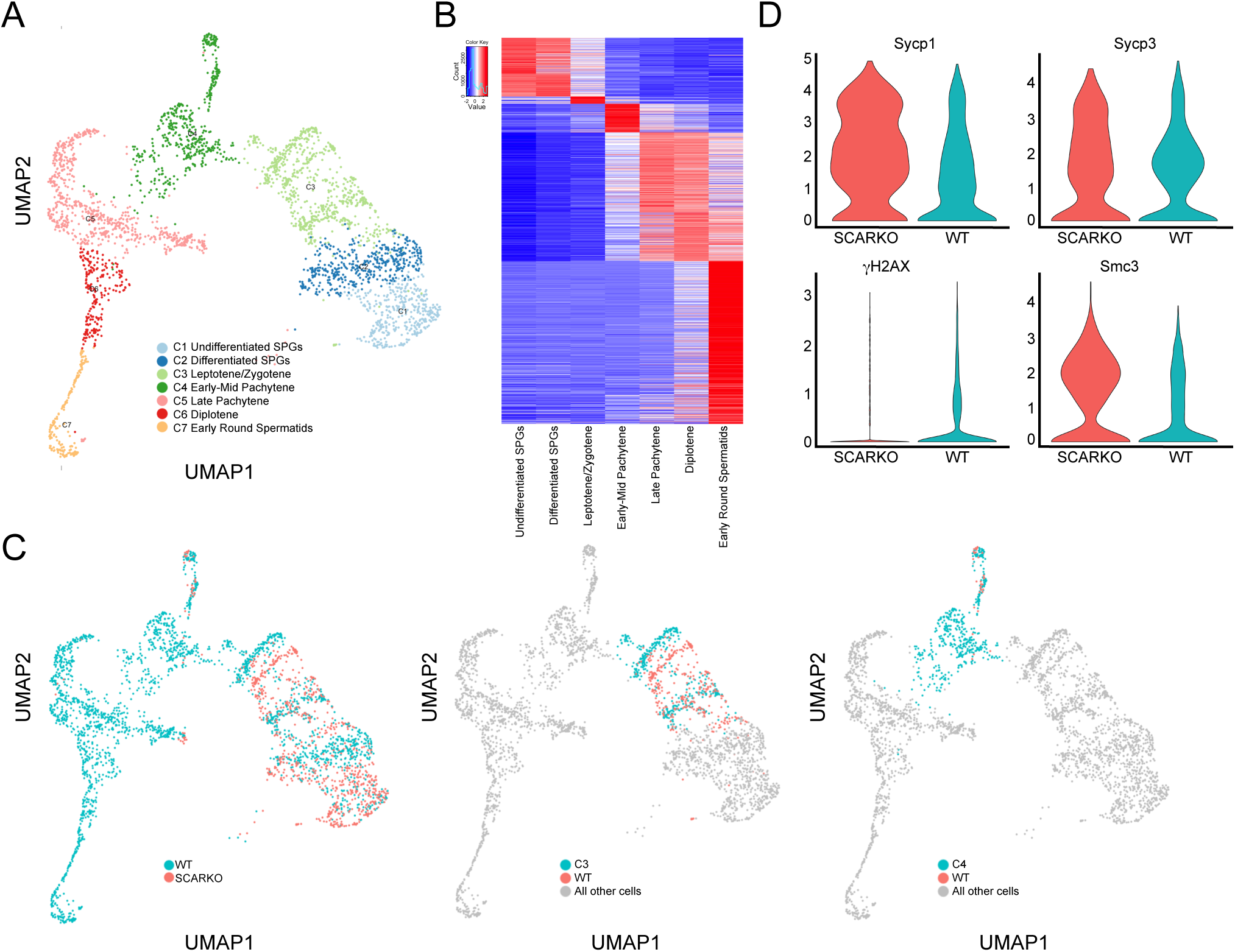
The SCARKO germ cell arrest occurs at the leptotene/zygotene to early-mid pachytene transition. A) Focused re-clustering of germ cells, regardless of genotype, identifies seven clusters. A UMAP plot of ∼3,100 germ cells with >500 detected genes, colored by assignment to 7 major clusters. B) Marker gene heatmap for the 7 clusters. Note – in the heatmap each marker gene is standardized over the 7 cluster centroids and ordered by cell type. C) Visualization of all germ cells post-quality control (QC) filter in UMAP overlaid by genotype contribution; both genotypes shown (left), cluster C3 containing leptotene/zygotene cells highlighting genotype contribution (middle) and cluster C4 containing early/mid-pachytene cells highlighting genotype contribution (left – pink). D) Representative violin plots for known meiosis markers within each genotype.

Our analysis of the germ cell trajectory **(Figure 5C**, left panel) revealed that the majority of SCARKO germ cells did not exhibit transcriptomic progress beyond the transition from leptotene/zygotene to early-mid pachytene stages (**Figure 5C**), with very few cells from SCARKO testes (32/800, 4%) reaching the early-mid pachytene transcriptional state compared to wild-type (461/2385, 19%). This observation was also reflected in differential expression of marker genes such as *Sycp1, Sycp*3, *H2afx*, and *Smc3* (**Figure 5D**). Hence, the scRNAseq analysis captures a transcriptome state in which SCARKO mutant germ cells have stalled in early pachytene, although they cytologically progress to a mid-pachytene-like state (**Figures 2, 3**). This implies that the molecular/transcriptomic state is uncoupled from the cytological progress of the germ cells in SCARKO mutant testes.

To identify genes that might underlie the germ cell arrest/loss in SCARKO mutants, we compared transcriptomes of C3 cells, which correspond to the stage prior to the arrest, between wild-type and SCARKO testes. We identified 25 genes that were at least 2-fold higher (p-value < 0.01) in wild-type germ cells **(Table 1**). Among these were genes previously known to be modulated by androgen signaling within the testes, such as *Fabp9* and *Gstm5* (De Gendt *et al.*, 2014). Other SCARKO downregulated genes included an RNA-binding protein (*Ybx3*), and proteins involved in cytoskeleton, acrosome or cilia organization and formation (e.g., *Meig1, Spink2*, and *Rsph1)*, which are important components for post-meiotic spermiogenic differentiation (Zhou *et al.*, 2010; Kott *et al.*, 2013; Snyder *et al.*, 2015; Zhang *et al.*, 2019). These observations suggest that expression of a core spermiogenesis gene program is required to transition from mid-meiotic to spermiogenic stages, which could imply either a permissive role, or an instructive, licensing role.

Given that germ-cell arrest is not fully penetrant, we next compared transcriptomes of SCARKO and WT germ cells that escaped the arrest (C4). Surprisingly, this analysis failed to identify any significant gene expression differences, suggesting that spermatocytes in SCARKO testes are transcriptionally similar to pachytene spermatocytes from wild-type testes. Indeed, these spermatocytes may represent the small population of spermatocytes from SCARKO testes that are competent to undergo cytological division-phase chromosome condensation after treatment with OA (**Figure 3G**).

### Loss of functional Sertoli androgen receptor alters the transcriptome of Sertoli cells but not of other testicular somatic cells

In addition to soma-to-germ or germ-to-soma communication, the intercellular communication within the testis somatic compartment is important for maintaining tissue homeostasis and function. Therefore, we next examined how loss of AR signaling from Sertoli cells may affect the transcriptome of Sertoli cells themselves or those of other somatic cells in the testis. The merged wild-type and SCARKO datasets consisted of 18,298 somatic cells: 12,373 in SCARKO and 5,830 in wild-type mice (**Figures 6A**). Higher relative abundance of somatic cells from the SCARKO mutants was expected, given the reduction in the germ cell populations. Clustering analysis of the ∼18K somatic cells revealed five clusters, which by marker gene analyses correspond to Sertoli cells, interstitial fibroblasts (*Tcf21*+), immature Leydig, adult Leydig, and macrophages **(Figure 6A**; **Supplemental Table S4).** We then compared the transcriptomes of wild-type and SCARKO mutants in each of the five somatic cell types (**Figure 6B**). However, we did not observe significant transcriptome differences in the somatic cell populations, namely macrophages, interstitial fibroblasts, immature and adult Leydig cell populations (data not shown). Next, the focused re-clustering of Sertoli cells only identified a single cluster of well-mingled WT and SCARKO mutant cells, suggesting that Sertoli cells from the two genetic backgrounds are transcriptionally similar (**Figure 6C**). This was further illustrated on a gene-by-gene level when we examined the average expression of ubiquitously expressed Sertoli cell markers, i.e. *Sox9* and *Rhox8*, between the two genotypes (**Figure 6D**). However, differential gene expression analysis identified a 115 genes with at least 1.5 fold difference (p-value < 1E-5) between wild-type and SCARKO Sertoli cells (**Table 2**). Sixty seven of these genes were downregulated in SCARKO Sertoli cells, 23 of which have been reported in the literature to be androgen-regulated, e.g. *Tsx, Susd3, Rhox5, Serpina5, Drd4, Eppin, Clstn1, Esyt3 and Jam3*, including *Ar* itself, as expected (Lindsey and Wilkinson, 1996; Tan *et al.*, 2005; Willems *et al.*, 2010; De Gendt *et al.*, 2014) (**Table 2**). Whereas, fourty eight genes were upregulated in SCARKO mutants. Therefore, loss of AR in Sertoli cells leads to misregulation of a small number of genes, which explains the lack of global transcriptome changes in the Sertoli cell population in SCARKO mutants. Taken together, our scRNA analysis revealed a discrete molecular fingerprint of AR-regulated genes in Sertoli cells, including known and previously unknown genes.

**Figure 6:**
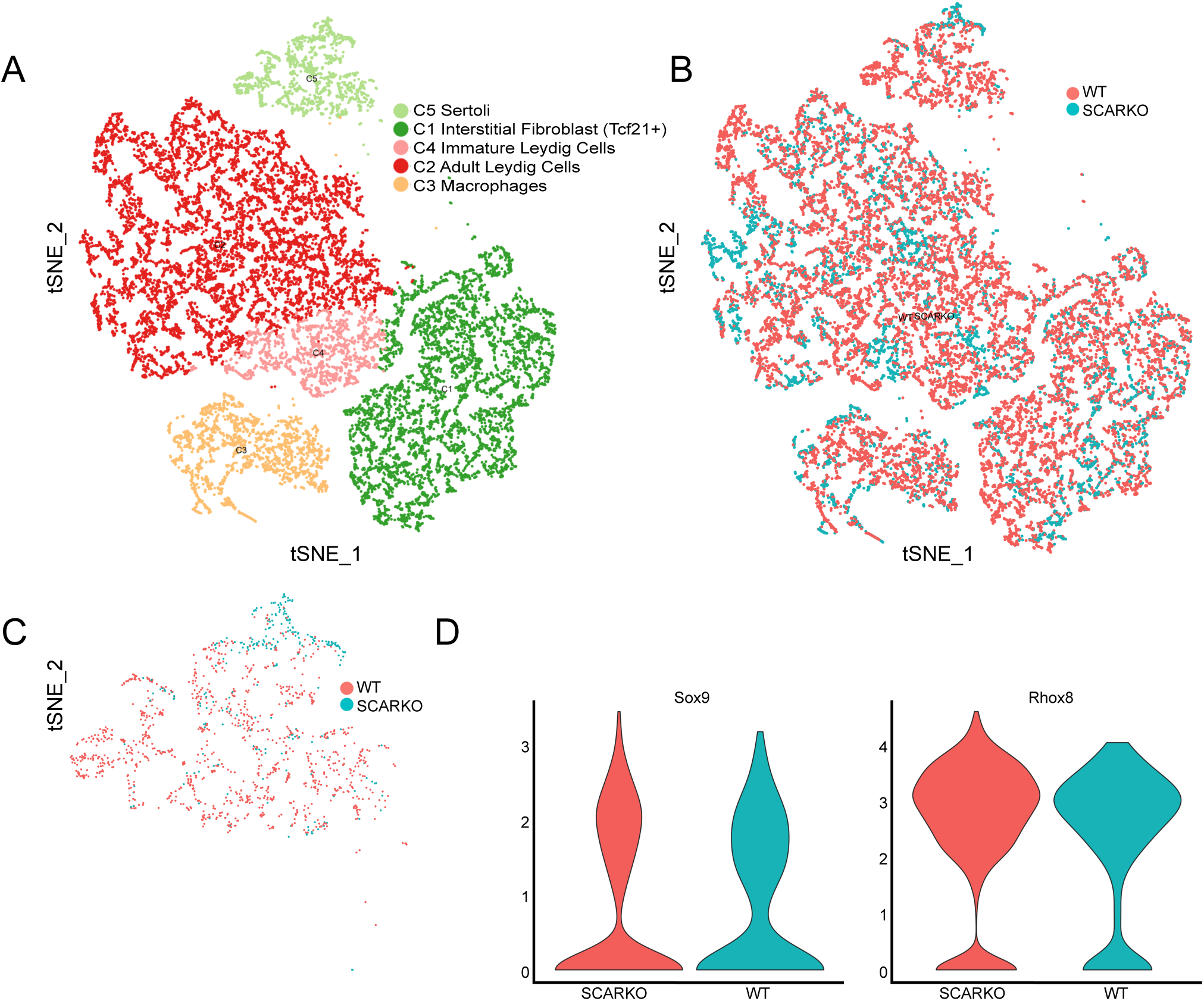
Loss of functional Sertoli androgen receptor alters the transcriptome of Sertoli cells but not of other testicular somatic cells. A) Focused re-clustering of 18,000 somatic cells identifies five major somatic cell types. TSNE projection of all somatic cells with >500 detected genes, colored by assignment to five major clusters. B) Visualization of all somatic cells post-quality control (QC) filter in TSNE overlaid by genotype contribution. C) Selection and Re-clustering of *Sox9+* and *Rhox8+* cells in TSNE space, colored by genotype, D) Violin plots illustrating expression levels of known Sertoli cell marker genes *Sox9* and *Rhox8* by genotype.

## Discussion

This study investigated the mechanistic basis upon which androgen signaling in Sertoli cells instructs progression of germ cells meiosis. Specifically, we exploited the SCARKO model coupled with synchronization of spermatogenesis in juvenile mice (**Figure 1; Supplemental Table 1**) and scRNAseq analyses of the juvenile testis. Our cytological analysis of SCARKO mutant testis shows extensive germ-cell loss at all stages of germ cell development (Figure 2), suggesting that AR signaling profoundly affects testicular homeostasis. Furthermore, we show that androgen signaling in Sertoli cells is not essential for major chromosomal events of meiotic prophase, including chromosome synapsis, reciprocal recombination, and expression of mid-pachytene markers that precede acquisition of competence for the meiotic division phase (Figure 3A-F). Despite an apparent normal meiotic progression, pachytene SCARKO spermatocytes fail to acquire competence to enter the division phase, and instead stall in a pachytene-like histological state (Figure 3G-H). Our single cell RNAseq confirmed the meiotic arrest phenotype, but also revealed a unexpected disconnect in germ cell cytological state vs. molecular states (Figure 4 and 5). Although the SCARKO mutant germ cells reach a pachytene like state histologically, they arrest at a leptotene or zygotene transcriptome state. Many of the genes that fail to activate in SCARKO mutant germ cells are involved in spermiogenesis, suggesting that AR-induced signaling in Sertoli provides an environment conducive for mid-prophase spermatocytes to undergo meiotic division and embark upon spermiogenic differentiation. In the somatic compartment of SCARKO mutants, we identified several AR regulated genes in Sertoli cells but did not observe any transcriptional changes in neighboring somatic cells. In all, our approach enabled us to uncover important germ cell intrinsic and extrinsic features required for proper germ cell progression.

### Loss of functional Sertoli androgen receptor alters the transcriptome of Sertoli cells

Our differential scRNAseq analysis identified 115 genes misregulaated in SCARKO Sertoli cells (Table 2), of those genes, 39 were previously reported by de Gendt et al., 2014, and previously associated with male infertility or subfertility when gene function is abrogated, suggesting biological significance for the set of 115 genes. For example, *Tsx* mutant mice are subfertile and have an increase in apoptotic pachytene cells (Anguera *et al.*, 2011) while *Bsg* knockout animals are sterile due to impaired interaction between germ cells and Sertoli cells (Bi *et al.*, 2013). In addition, *Serpina5* mutant mice exhibit compromised blood-testis barrier function (Uhrin *et al.*, 2000), a phenomenon also noted in SCARKO mutants. Given that many of the 115 genes lead to infertility in a loss of function context, we hypothesize that gene combinations or epistatic interactions may contribute to diminished function of SCARKO Sertoli cells and infertility in SCARKO animals. Interestingly, no transcriptional differences were observed in the neighboring somatic cell types. This suggests that loss of androgen signaling within juvenile Sertoli cells is not immediately impacting the environment surrounding the seminiferous tubules. This observation is consistent with an earlier report in which loss of AR signaling in the testis didn’t result in altered Leydig cell number in the juvenile testis (O’Shaughnessy *et al.*, 2002). However, this study found changed Leydig cell numbers in aging mice, suggesting that transcriptional alterations in extra-tubular somatic populations develops after subsequent rounds of spermatogenesis and onset of puberty.

### Sertoli-cell androgen signaling is required for the normal germ-cell transcriptome and for onset of meiotic divisions

In spite of the fact that male germ cells lack cell-autonomous AR expression, a large body of evidence demonstrates that androgen signaling from surrounding Sertoli cells is required for normal progress through spermatogenesis (O’Hara and Smith, 2015). In addition to the known spermatocyte meiotic arrest in SCARKO testes, we show here that germ cells undergo pervasive apoptosis throughout meiotic prophase. This data show that androgen signaling from Sertoli cells throughout meiotic prophase is critical, most probably by providing a nurturing environment for germ cells. Importantly, however, a subset of spermatocytes survive and progress normally through central meiotic processes, undergoing chromosome pairing and synapsis, XY body formation, and and expressing markers of meiotic recombination (Figure 3). Hence, quintessential prophase chromosomal dynamics, defining events from leptotene to mid-pachytene, can occur independently of androgen signaling from Sertoli cells. However, despite an apparent normal chromosomal progression, SCARKO spermatocytes are not competent to undergo the natural transition from meiotic prophase to the division phase, nor are they competent to respond to a pharmacological agent that promotes premature transition from meiotic prophase (Figure 3H). Together, the phenotype of normal cytological features of mid-pachynema combined with lack of competence to progress to the division phase suggests that androgen signaling is required for germ cells to acquire competence for meiotic division and subsequent spermiogenesis.

Very little is known about spermatocyte gene expression required for meiotic division-phase competence, an issue directly tackled by this study. At a general level, it is known that gene expression changes during meiotic prophase are extensive (De Gendt *et al.*, 2014; Chen *et al.*, 2018; Green *et al.*, 2018; Jung *et al.*, 2019), with modules or clusters of genes expressed at appropriate times for ongoing prophase events (Jung *et al.*, 2019). Nonetheless, genes known to be required for fertility tend to reach peak expression levels in a cell stage at or preceding the stage of observed cytological arrest (Green *et al.*, 2018). Moreover, bulk tissue sample analysis also reveal that some transcripts are expressed in prophase spermatocytes but are not required or translated until post-meiotic spermiogenesis (Ball *et al.*, 2016). Our goal here was to define the androgen-dependent molecular signature and the precise germ cell stage of arrest. Our cytological and scRNA-seq analyses showed that the majority of SCARKO germ cells experienced a transcriptomic stall point at the leptotene/zygotene transition, but progressed further based cytological assesment. The transcriptomic arrest of most SCARKO spermatocytes at a leptotene-zygotene state raised the questions not only of function of genes failing to be expressed, but also whether there might be aberrant gene expression at even earlier stages of spermatogenesis. In fact, some of the genes identified as differentially expressed in the leptotene/zygotene mutant transcriptomes were expressed earlier (Table 1), including three as early as in undifferentiated spermatogonia (*Ldhc, Meig1* and *Fabp9*) and 16 in differentiating spermatogonia. The remaining mis-regulated genes were specifically expressed in leptotene/zygotene (*Crisp2, Meig1, Spink2* and *Gstm5*; Table 1). Some of the identified genes are required for organization of the cytoskeleton, acrosome or cilia, structures that are formed in later germ cells; therefore, it appears that transcripts required for meiotic and spermiogenic competence are accumulated prior to the time of their developmental need.

In the absence of a functional Sertoli cell androgen receptor, germ cells progressively acquire aberrant transcriptomic attributes (a germ-cell “SCARKO molecular phenotype”), culminating in a terminal leptotene-zygotene transcriptional signature with failure to progress to subsequent transcriptomic signatures. In spite of this pervasive transcriptomic block, a few (∼4% of the sequenced germ cells) mutant spermatocytes are found in the early-mid pachytene cluster, and are indistinguishable at a transcriptional level from the similarly staged wild-type spermatocytes. The small subset of cells that escape the leptotene/zygotene transcriptional block may correspond to the rare spermatocytes in the SCARKO testes that are capable of condensing chromosomes in response to OA treatment *in vitro (Figurre 3G-H)*.

Our findings highlights a discrepancy between a seemingly normal meiotic progress of surviving spermatocytes to the mid-pachytene stage, as observed with cytology analysis, and an apparent transcriptional arrest for most cells at an earlier stage (comparable to the WT transcriptome of leptotene-zygotene spermatocytes). Thus it should not be assumed that cytological and/or morphological states are necessarily coupled with corresponding molecular transcriptomic states. Indeed, a similar transcriptional uncoupling effect was recently found by studying a *Prdm9* mutation (Fine *et al.*, 2019), a gene required for selective activation of hotspots of meiotic recombination. In this report, *Prdm9* mutant germ cells exhibited cytological arrest at a pre-pachytene stage, but RNA-seq analyses revealed expression of genes that are normally activated later in meiotic prophase that support subsequent spermiogenesis. Together, these findings suggest that the temporal pace of gene activation and expression is not necessarily tied to the temporal events of meiotic prophase chromosomal dynamics.

In conclusion, synchronization of spermatogenesis coupled with single cell RNAseq analysis of SCARKO and aged matched wild type testes enabled us to identify and molecularly dissect the precise time point of androgen-dependent germ cell arrest. Our observations suggest that the Sertoli cells themselves create a permissive environment for meiosis and subsequent spermiogenesis. Future studies stand to discover unknown mechanism underlying critical meiotic prophase transitions in spermatogenisis by investigating the specific genes identified here.

## Acknowledgements

We thank members of the Hammoud, LI and Handel lab for scientific discussions and manuscript comments. We acknowledge the able assistance of Catherine Brunton for maintenance of mice and Karen Davis for assistance in preparing figures. This research was supported by National Institute of Health (NIH) grants 1R21HD090371-01A1 (S.S.H.), 1DP2HD091949-01 (S.S.H.), and Michigan Institute for Data Science (MIDAS) grant for Health Sciences Challenge Award (S.S.H.), CTRB training grant 5T32HD079342-04 (A.N.S.), MSTP training grant 5T32GM007863-38 (A.N.S.).

## Materials and Methods

### Animals

All experiments were conducted following Institutional Animal Care and Use Committee (IACUC) at The Jackson Laboratory and University of Michigan. All mice used in our experiments were obtained from a mixed genetic background C57BL/6; 129/SvEv. Specifically, we crossed the *Ar* ^fl/fl^ females (Chakraborty *et al.*, 2014)with Amh-cre ^tg/+^ (Lécureuil *et al.*, 2002); yielding ∼50% of the resulting males lacking AR in Sertoli cells (hereafter referred to as SCARKO). The Amh-Cre^-^ male littermates were used as wild type (WT) controls. Animal genotypes were confirmed by PCR as previously described (Chang *et al.*, 2004).

### Synchronization of seminiferous epithelium

Onset of spermatogenesis was synchronized as previously described (Hogarth *et al.*, 2013). Briefly, WIN 18,446 was administered to pups orally from 2-8 dpp at a dose of approximately 100 mg/kg/day. At 9 dpp, pups were given an IP injection of RA (100ug/animal). Testes were collected between 6 to 16 days following the RA injection (**Table 1**).

### Meiotic chromosome spreads

Meiotic spreads were prepared as previously described (Peters *et al.*, 1997). Briefly, testes were harvested, detunicated, and then incubated for 45 min in hypotonic-extraction buffer (30mM Tris-HCl, 50mM sucrose, 17mM sodium citrate, 5mM EDTA, 2.5mM DTT, 0.5mM PMSF, and 1X protease inhibitor). Cells were mixed in 100mM sucrose and spread on slides that were pre-washed with 0.5% PFA/0.05% Triton X-100. Slides were kept in humid chamber approximately 16hrs at 4°C. Slides were then air dried, washed in 0.04 % Photo-Flo 200 solution (Electron Microscopy Sciences) in ddH2O for 1hr, and air dried completely. The slides were then either stained immediately or stored at -20°C for later use. Immunofluorescence

For immunostaining chromosome spreads, slides were blocked for 1h at room temperature in blocking buffer (0.3 % Bovine Serum Albumin (BSA), 1% goat serum in phosphate-buffered saline with 0.05% Triton X-100). The slides were incubated with primary antibodies approximately 16hrs at 4°C. Immunolabeling was done using the following primary antibodies: Guinea pig anti-H1t (1:100, Handel Laboratory), mouse anti-γH2AX (1:100, Invitrogen Cat# MA1-2022), mouse anti-MLH1 (1:20, Abcam Cat# ab14206), rabbit anti-SYCP1 (1:100, Novus Biologicals Cat# nb300-229), rabbit anti-SYCP3 (1:100, Novus Biologicals Cat# nb300-232), rat anti-SYCP3 (1:100, Handel Laboratory). Slides were washed with the blocking buffer and incubated for 4 hrs with conjugated secondary antibodies and counterstained with DAPI to identify nuclei. Secondary antibodies conjugated with either Alexa 488 or 594 were used for these analyses (Thermo Fisher Scientific). Cells were visualized and quantified using a microscope (Leica Leitz DMRD).

### TUNEL staining

Testes were collected, fixed in 4% PFA (in PBS) for approximately 16hrs at 4°C, dehydrated in ethanol wash series, and embedded in paraffin. Four-micron FFPE tissue sections were deparaffinized, rehydrated, and boiled in 0.01M sodium citrate, pH 6.0, for 3min. After antigen retrieval, tissue sections were blocked in 20% horse serum, 0.3% BSA in PBS for 30min at room temperature and processed following the Roche Applied TUNEL kit. All images collected used a Leica Leitz DMRD microscope. The percent TUNEL positive cells was determined by averaging at least 100 tubules for each sample.

### In vitro induction of metaphase

This procedure was adapted from several sources (Evans *et al.*, 1964)(Wiltshire *et al.*, 1995; Cobb *et al.*, 1999). Briefly, testes were dissociated using 0.5 mg/ml collagenase diluted DMEM + 25mM HEPES for 20 min at 37°C with shaking, washed three times in DMEM, then digested with 0.5 mg/ml trypsin for 12 min at 37°C with shaking to generate a single cell suspension. The mixed germ cell suspension were filtered through a 70 micron Nitex mesh and washed 3 times in DMEM + 25mM HEPES medium. Approximately 100,000 cells were placed into culture dish and treated with either 400µM OA stock solution or vehicle (100% ethanol), respectively for 4 hours. The cultured spermatocytes were collected through centrifugation and washed in 2.2% sodium citrate, then 1% sodium citrate. The cells were then fixed (3:1 ratio of 100% ethanol; glacial acetic acid with 2% chloroform) for 5min at room temperature with gentle rocking. Fixed germ cells were then dropped from a height of approximately 0.5m on a glass slide and were allowed to air dry before being stained with Giemsa for the presence of condensed metaphase cells.

### Isolation of single cells for sequencing

Testes from transgenic SCARKO mice and WT litter mates were transferred to 10ml of digestion buffer 1 (comprised of Advanced DMEM:F12 media (ThermoFisher Scientific), 200 mg/ml Collagenase IV (Sigma), and 400 units/ml DNase I (Sigma). Tubules were dispersed by gently shaking by hand and allowed to settle for 1 min at room temperature. The supernatant was collected, placed on ice and quenched with the addition of fetal bovine serum (FBS) (ThermoFisher Scientific). The remaining tubules were then transferred to digestion buffer 2 (0.25 mg/ml trypsin (ThermoFisher Scientific) and 400 units/ml DNase I (Sigma) dissolved in Advanced DMEM:F12 media) and dissociated at 35C / 215 rpm for 5 minutes each. Any remaining tubule fragments were broken up with manual agitation using a 1000uL pipette. The resulting single cell suspension was transferred to the previously collected supernatant and quenched with the addition of fetal bovine serum (FBS) (ThermoFisher Scientific). Cells were filtered through a 100 µm strainer, washed in Phosphate-buffered saline (PBS), pelleted at 300g for 5 minutes, and re-suspended in MACS buffer containing 0.5% BSA (Miltenyi Biotec). For all Drop-seq experiments, the live single-cell suspensions were collected by flow cytometry using (FACS) ARIA II/III (BD Biosciences) cell sorter.

### Drop-Seq Procedure

Single-cell suspensions were diluted to 250 cells/ml and processed as described previously (Macosko *et al.*, 2015; Green *et al.*, 2018). Briefly, cells, barcoded microparticle beads (MACOSKO-2011-1-0, Lots 072817 and 060718, ChemGenes Corporation), and lysis buffer were co-flown into a microfluidic device and captured in nanoliter-sized droplets. After droplet collection and breakage, the beads were washed, and cDNA synthesis occurred on the bead using Maxima H-minus RT (Thermo Fisher Scientific) and the Template Switch Oligo. Excess oligos were removed by exonuclease I digestion (New England Biosciences). cDNA amplification was done for 15 cycles from pools of 2,000 beads using HotStart ReadyMix (Kapa Biosystems) and the SMART PCR primer. Individual PCRs were purified and pooled for library generation. A total of 600 pg of amplified cDNA was used for a Nextera XT library preparation (Illumina) with the New-P5-SMART PCR hybrid oligo, and a modified P7 Nextera oligo with 10 bp barcodes. Sequencing was performed on a HiSeq-4000 (Illumina) 51 x 151nt with the Read1CustomSeqB primer.

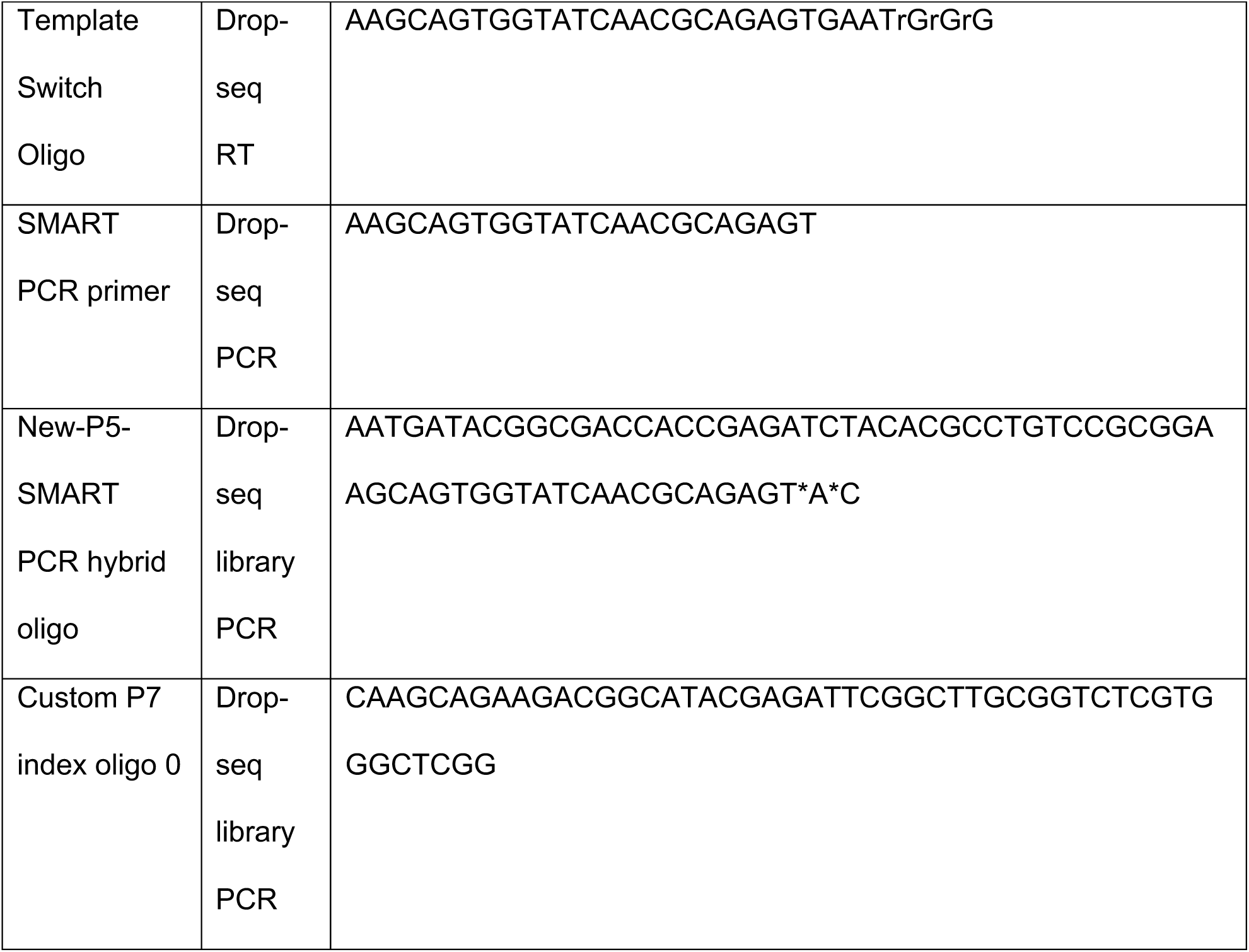
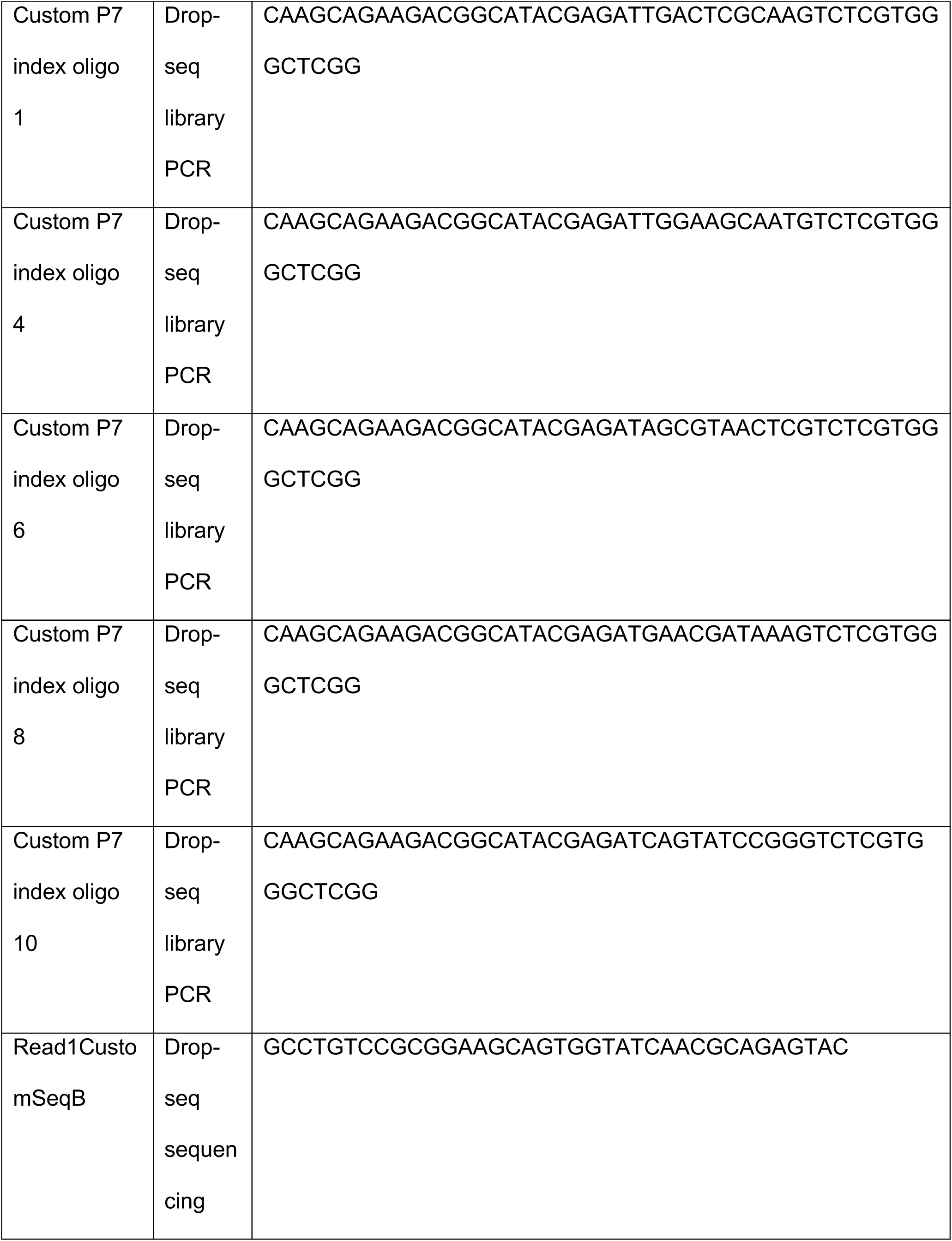

### Computational analysis of Drop-Seq Data

#### CCA batch correction and clustering all genotypes and time points to identify germ cells and interstitial cells

All samples were aligned using the canonical correlation analysis (CCA) embedded in the R package Seurat v2.3.4. Cells with < 500 detected genes or with >10% of transcripts corresponding to mitochondria-encoded genes were removed, as previously described (Green *et al.*, 2018). The CCA analysis was performed by taking the union of the top 2,000 highly variable genes. The top 10 significant CCs - determined by MetageneBicorPlot - were used for Louvain-Jaccard clustering, resulting in seven clusters with a resolution of 0.3. Markers for each of the clusters were obtained by comparing a given cluster with all other clusters using the binomial likelihood test embedded in R package Seurat. Marker Selection criteria: (1) genes with a minimum difference in the fraction of detection between the two groups of at least 20%; (2) a minimum of 2-fold enrichment in the cluster compared to all other clusters, and (3) a binomial corrected p-value < 0.01.

#### Evaluation of between-batch reproducibility

Each sample was subset from the overall CCA alignment, and for each batch, we calculated cluster centroids of each gene as ln(mean of normalized expression+1) over all cells in each cluster. To compare the cluster centroids among the eight batches, we cross-tabulated rank correlation coefficients among all pairs of cluster centroids across the eight samples, which demonstrated that clusters were largely reproducible across batches.

#### Identifying and merging germ cells from all genotypes and time points

For each individual time point, the WT and SCARKO mutant were aligned using the CCA and clustered by aligned CCs as described above. Germ cell clusters were defined for each individual time point by evaluating the markers and used to filter the raw data matrix to contain only germ cells. We then merged the raw data matrices from the germ cell subset of all samples and performed principal component analysis (PCA) on the standardized gene expression matrix. To assess cellular heterogeneity, we performed Louvain-Jaccard clustering to identify cell clusters using the top eight PCs (Blondel *et al.*, 2008). After clustering, we repeated calling markers following the selection criteria above to identify the subtypes of germ cells present within the merged object.

#### Differential expression across germ cell clusters

To identify genes dynamically regulated between clusters, we called differentially expressed genes between the two clusters regardless of time point or genotype using the binomial likelihood test embedded in R package Seurat with a minimum of 2 fold-change and p-value < 0.01. To identify genes differentially expressed between two genotypes (WT and SCARKO) within each germ cell development stage, we again called differentially expressed markers using the same parameters as above, but between the two genotypes within a cluster.

#### Sertoli cell analysis

From the overall CCA alignment of all cells, we defined the Sertoli cell cluster using top markers. The identified Sertoli cell cluster was subset from the rest of the interstitial and germ cells. These cells were then visualized in t-SNE space. Differential gene expression between WT and SCARKO genotypes was then calculated using the using the binomial likelihood test with a minimum of 1.5 fold-change and p-value < 0.01. As a result, we obtained 30 markers that are expressed high in WT but mis-regulated in SCARKO. To examine if these 30 mis-regulated markers are specific for any of the 9 Sertoli stages previously identified from normal adult mouse testis, we calculated the “mis-score” for each of Sertoli cells in adult mouse testis as the total expression of these 30 mis-regulated genes divided by the total expression of all detected genes, and then visualized the “mis-score” for each of the 9 sertoli stages in Violin plot.

---

**Figure S1:**
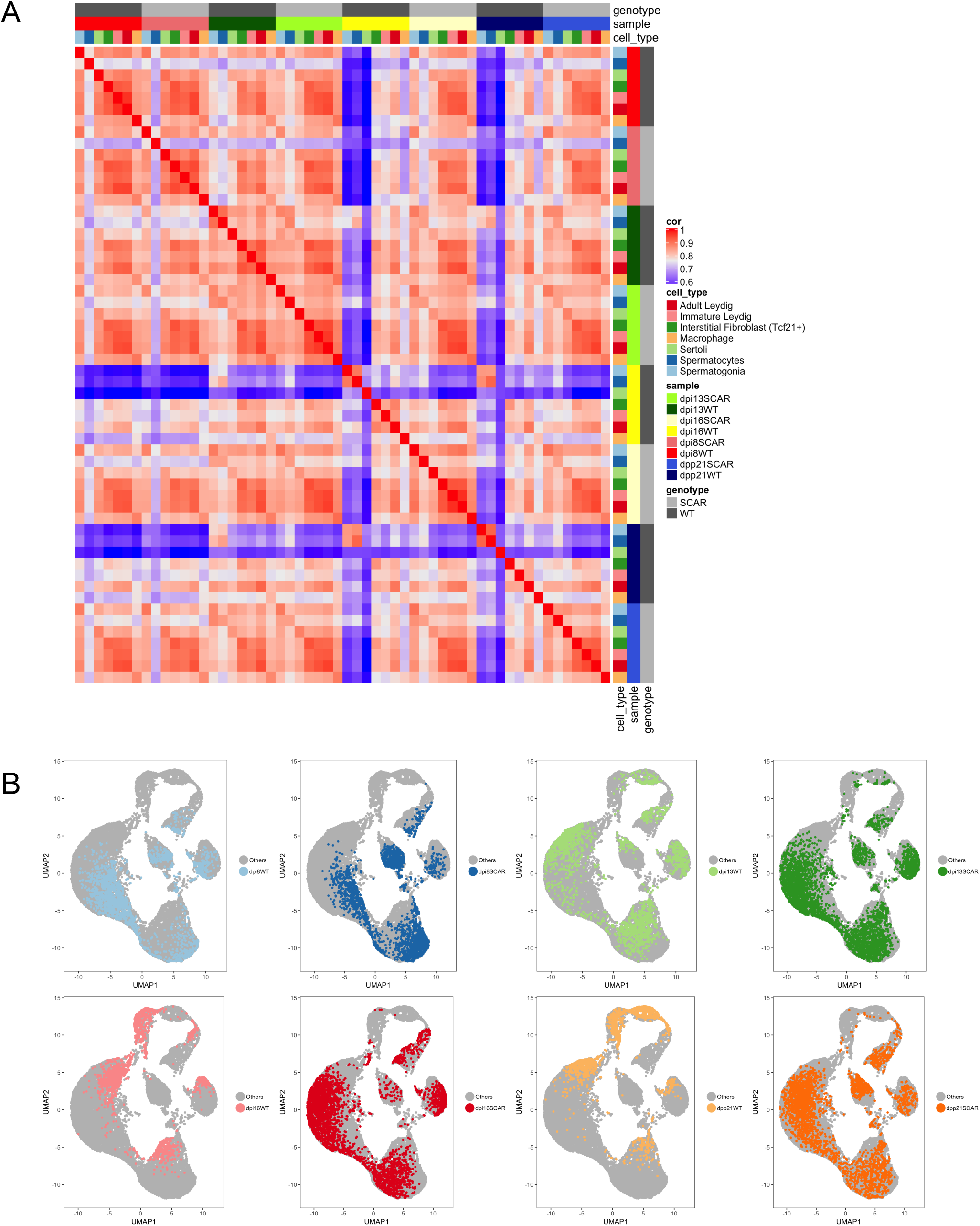
Evaluation of sample reproducibility across both genotypes and all time points analyzed. A) Heatmap of pairwise rank correlation matrix among the 8 sample cluster centroids for each of the 7 cell-types. Bars on right and top demarcate either genotype, sample time point or cell types. B) Contribution of each sample to overall CCA alignment of all 26,500 cells, visualized in UMAP space.

## Notes

Support : This work was supported by NIH HD33816 (M.A.H), 1R21HD090371-01A1 (S.S.H. and J.Z.L.), 1DP2HD091949-01 (S.S.H.), F30HD097961 (A.N.S) and training grants 5T32HD079342 (A.N.S.), 5T32GM007863 (A.N.S.), Michigan Institute for Data Science (MIDAS) grant for Health Sciences Challenge Award (J.Z.L. and S.S.H.), Open Philanthropy Grant 2019-199327 (5384, S.S.H.), and partially supported by NIH P30 CA034196 to the Jackson Laboratory. The content is solely the responsibility of the authors and does not necessarily represent the official views of the NIH.

### Competing Interest Statement

The authors have declared no competing interest.

